# mehari: high-performance, strict HGVS-first variant effect prediction

**DOI:** 10.64898/2026.05.12.724271

**Authors:** Till Hartmann, Max Zhao, Dieter Beule, Manuel Holtgrewe

## Abstract

Variant annotation requires the precise and consistent computation of Sequence Ontology (SO) terms and Human Genome Variation Society (HGVS) nomenclature. To ensure robust synchronization between these two key facets, we present mehari, a high-performance variant effect predictor implemented in Rust that employs a strict “HGVS-first” approach. By deterministically projecting variants to transcripts before evaluating functional consequences, mehari structurally aligns HGVS notation and SO terms. Benchmarking on ClinVar demonstrates that mehari achieves exceptional processing speeds and high concordance with established tools like Ensembl VEP, while also providing refined handling for complex biological edge cases such as selenoprotein recoding.

## Introduction

High-throughput sequencing for genomic variant detection has become the backbone of clinical genomics. After the variant calling step, annotating the variant is a critical step for clinical genomics as it is a requirement for their interpretation. The annotation of variants is a broad field ranging from clinical/population database lookups, e.g., gnomAD, ClinVar, or dbSNP to computationally expensive steps such as deep learning–based prediction on splicing with SpliceAI or protein folding with AlphaFold.

The annotation with so-called functional consequence (missense, frame-shift, etc.) is at the core of this annotation process. Based on genome annotation with genes’ transcript models, this step computes the molecular impact for each variant and each overlapping transcript. This annotation has two facets: annotation with Sequence Ontology (SO) terms and with Human Genome Variation Society (HGVS) syntax on the transcript and most relevantly on the protein level. Both facets aim to standardize variant descriptions: the HGVS notation provides an unambiguous coordinate and sequence description, while the Sequence Ontology provides a standardized vocabulary for describing variation. Crucially, standardizing on official SO terms avoids the introduction of tool-specific definitions.

Such annotations are challenging as they must be both correct (assuming the correctness of transcript models) and fast (given the rapidly growing nature of genomic datasets). There is neither room for ambiguity (as the HGVS nomenclature is well-defined and journals require validation by the “blessed” tool VariantValidator.org) nor for waste of resources. Additionally, comprehensive tools must support diverse transcript databases like Ensembl and RefSeq. RefSeq models in particular introduce additional complexity, as their transcript-to-genome alignments can contain indels that must be modeled correctly during coordinate projection.

One issue with legacy annotation tools, such as `VEP`, is the potential for structural mismatches between SO and HGVS, as they are computed independently. While the aim of the HGVS notation is the unambiguous description of variants, the description of SO leaves room for ambiguity as there is no equivalent of the 3′ rule per se. Such inconsistencies unnecessarily complicate downstream clinical interpretation and can lead to inaccurate clinical representations.

To address this, we introduce `mehari`, a **fast and correct** variant effect predictor, implemented in the Rust programming language. Rather than emulating legacy idiosyncrasies, `mehari` is a strict and rigorous predictor utilizing an “HGVS-first” approach. Given a VCF file, variants are interpreted as HGVS genomic variants, projected and normalized to transcripts and finally to proteins. This methodology guarantees that the variant notation and the functional prediction remain perfectly synchronized.

## Materials and Methods

### System Design & Architecture

`mehari` is written in the Rust programming language with the aim of correctness, speed, and memory safety. We implemented core functionality of variant representation (and parsing) in the re-useable library `hgvs-rs`. This library is a near 1:1 Rust port of the established Python `biocommons-hgvs` package which also powers the *de facto* standard in variant representation: VariantValidator.org. `hgvs-rs` handles the step-by-step projection of variants from genomic (`hgvs.g`) to coding (`hgvs.c`) and protein (`hgvs.p`) coordinates. This process strictly enforces normalization rules, such as 3-prime shifting, before `mehari` evaluates any functional consequences.

To perform these projections, `mehari` takes an input variant and searches for all overlapping transcripts by querying an array-backed interval tree (provided by `rust-bio`). A default padding of 5kbp is added to capture up- and downstream variants. Once the coordinate projections are complete, `mehari` infers the Sequence Ontology (SO) terms. The level of inference depends on the functional impact. Certain terms are derived from exon alignments, while others are evaluated at the `hgvs.c` or `hgvs.p` level. The logic follows standard SO definitions very closely. It applies special handling only for specific biological edge cases. Finally, the tool formats and writes the results into a VCF or BCF file.

`mehari` is designed for versatile integration. It can be used both via a standard command-line interface or using an integrated REST API server. The server is capable of asynchronous execution using the `actix` library and provides an OpenAPI schema via `utoipa`. Additionally, it serves as a Python library via `pyO3`. The Python bindings provided via the `pyO3` library support SPDI string inputs, `kwargs`, and direct integrations with Polars DataFrames and Apache Arrow. Beyond transcript models, `mehari` supports auxiliary RocksDB databases for ClinVar and gnomAD annotations.

### Transcript Database Management

`mehari` builds its custom transcript model database either from cdot JSON files (originally derived from official Ensembl and RefSeq GFF/ GTF files) or from standard GFF3 files. The build snake-make[1] workflow takes the cdot JSON file and a SeqRepo instance containing the cDNA sequences. It constructs a custom transcript model from these inputs, performs sanity checks, and writes the results to a zstd-compressed protobuf representation. The complete workflow is available at https://github.com/varfish-org/mehari-data-tx. A major advantage of this approach is the exceptionally small storage footprint. As shown in Table 1, the `mehari` transcript database requires about 166MB, compared to `vep`’s roughly 24GB cache.^1^

**Table 1:** Data sources utilized by benchmarked annotators.

| Tool | Version | Database | Sources | Size (MiB) |
| --- | --- | --- | --- | --- |
| annovar | 2022-08-02 | hg38_refGeneWithVer | RefSeq | 316 |
| jannovar | 0.36 | refseq_curated_109_hg38 | RefSeq | 31 |
| mehari | 0.41.2 | mehari-data-tx 0.12.0 | Ensembl 115, RefSeq RS_2025_08 | 166 |
| nirvana | 3.18.1 | 91.27.66 | Ensembl, RefSeq | 1441 |
| snpEff | 5.4.0c | GRCh38.p14 | RefSeq | 637 |
| vep | 115 | Ensembl 115 | Ensembl 115 | 24 161 |

**Table 2:** Performance Benchmark.

| Tool | # annotated transcripts | CPU time (s) | CPU time / transcript (μs) |
| --- | --- | --- | --- |
| annovar | 3 572 463 | 993 | 277.95 |
| jannovar | 9 399 143 | **192** | 20.42 |
| mehari | **110 948 714** | 561 | 5.05 |
| nirvana | 40 008 357 | 195 | **4.87** |
| snpEff | 38 852 157 | 992 | 25.53 |
| vep | 74 668 641 | 21 791 | 291.83 |

### Benchmark Data Sources

We use the ClinVar GRCh38 dataset from ClinVar 2025-08-06 to evaluate `mehari`. This dataset is used both to compare annotations against `vep`[2] and to benchmark performance against `annovar`[3], `jannovar`, `nirvana`[4], `snpEff`[5], and `vep`. We restrict the ClinVar dataset to autosomal, gonosomal, and mitochondrial chromosomes. We also filter out variants with multiple alternative alleles. This leaves 3 572 463 variants for the evaluation.

**Figure 1:**
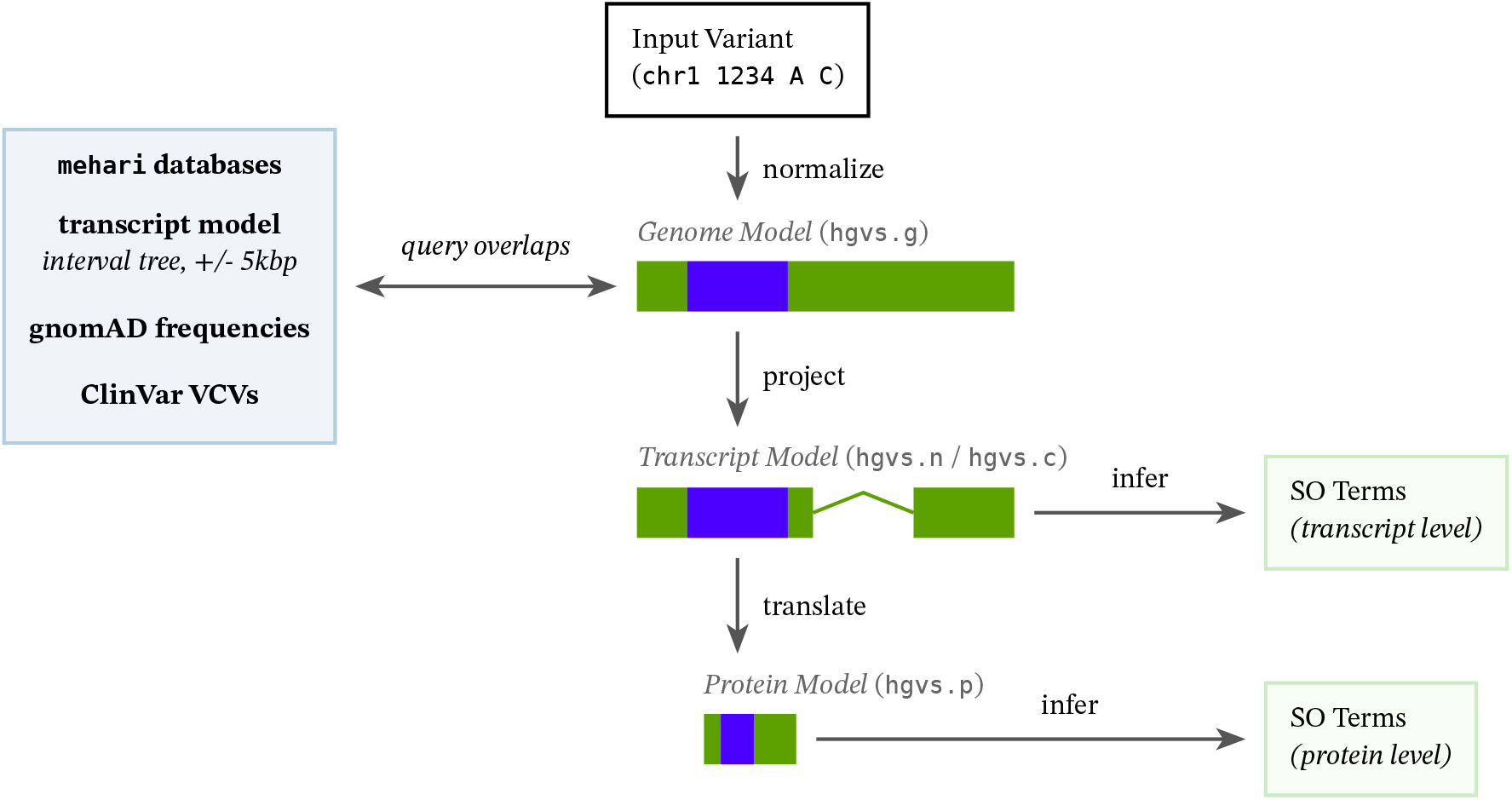
Architecture of `mehari`’s “HGVS-first” variant effect prediction. By projecting coordinates first, and mapping them to biological models, HGVS notation and Sequence Ontology functional predictions remain strictly synchronized.

For the annotation comparison, both `mehari` and `vep` use Ensembl transcripts from version 115 for GRCh38. For the performance benchmarks, we note that tools rely on different data sources and custom databases. They also have differing levels of up-to-date-ness, as detailed in Table 1.

### Evaluation & Benchmarking Framework

The workflow for benchmarking performance is available at https://github.com/varfish-org/mehari-benchmark. It runs the different tools multiple times and collects runtime statistics. By default, the benchmark uses exactly 1 core and 1 job at a time. This constraint is designed strictly to measure baseline algorithmic efficiency per transcript, rather than native parallelization limits.

The workflow for comparing annotations between `mehari` and `vep` is available at https://github.com/varfish-org/mehari-annotation-comparison. Because `vep` and `mehari` use different levels of specificity for SO terms, we employ two methods to normalize the output for comparison: First, we run `mehari` with a `--vep-compatibility-mode`. This mode heuristically recreates some of `vep`’s idiosyncrasies and adjusts for consequence terms `vep` does not use (e.g., replacing `non_coding_transcript_intron_variant` with the broader `intron_variant`, or altering the handling of `stop_gained` during frameshifts). Second, we filter the outputs of both tools using `vembrane`[6] to retain only the `most_specific_terms()` based on the SO directed acyclic graph. This prevents flagging trivial hierarchical discrepancies (like `intron_variant` vs `coding_transcript_intron_variant`) as hard differences, allowing us to isolate true biological discordance. Finally, the workflow utilizes external parallelization (“scatter-gather”) to annotate the ClinVar VCF in 24 parts.

## Results

### Performance Benchmarking

We ran `annovar`, `jannovar`, `mehari`, `nirvana`, `snpEff`, and `vep` on the ClinVar dataset. To ensure accuracy regardless of system load, we measured CPU time rather than wall-clock time. All of these tools rely on their own databases, utilizing different data sources in different versions (see Table 1). Because of these differences, the number of annotated transcripts varies between the tools. This directly influences both runtime and I/O. To provide a reasonably fair comparison, we normalize CPU time by the total number of annotated transcripts.

While `mehari` is not the fastest of all evaluated tools, it is the fastest classical VCF-in/VCF-out tool. Its processing speed per transcript is very close to `nirvana`. However, it should be noted that direct performance comparisons have inherent limitations: For one, `nirvana` does not produce VCF output, instead outputting JSON. Because downstream clinical tools almost universally expect VCF formatting, this renders `nirvana` less attractive as a direct architectural equivalent. Additionally, `nirvana` automatically annotates variants with allele frequencies from dbSNP and gnomAD by default, introducing additional I/ O and processing overhead that is not present in a baseline transcript-only annotation run.

### Sequence Ontology Term Concordance

`mehari` and `vep` annotations **completely** agree on 99.93 % of ClinVar variants. There are 2489 variants where `mehari` and `vep` produce differing annotations. For this comparison, `mehari` was run with its `--vep-compatibility-mode`.^2^ Conversely, `vep` was run with the `--shift_3prime 1` flag.^3^

To ensure a robust comparison of systematic differences rather than noise, we filtered the outputs of both tools using `vembrane`. This step retained only the `most_specific_terms()` based on the SO graph. For example, it filters out the broad parent term `intron_variant` if the specific child term `splice_acceptor_variant` is present. Grouping the resulting discordant annotations helps identify distinct categories of disagreement. These categories range from explicit tool artifacts to the limitations of `mehari`’s compatibility mode in mirroring `vep`’s internal logic, see Table 3.

**Table 3:** Summary of discordance patterns between vep and mehari consequence predictions on ClinVar variants. Counts reflect the occurrences of specific discrepancies; because a single variant may exhibit multiple artifacts simultaneously, categories are not mutually exclusive.

| Category | Count | Explanation |
| --- | --- | --- |
| Misplaced start_retained_variant | 809 | vep annotates start_retained_variant for variants far downstream of the canonical start. |
| Structural vs. Functional | 497 | While mehari's --vep-compatibility-mode generally suppresses stop_gained when a frameshift occurs to match vep, this can in some cases lead to discrepancies where vep *does* call stop_gained correctly. |
| Splice Term Hierarchy | 458 | For variants spanning adjacent functional regions (e.g., an acceptor site and a polypyrimidine tract), vep independently triggers and reports both. mehari's --vep-compatibility-mode explicitly suppresses the lower-impact tract term to enforce a strict regional hierarchy. |
| Missing High-Impact Terms | 220 | For large or complex deletions spanning multiple functional regions (e.g., UTR through CDS), vep frequently drops specific high-impact terms like start_lost or stop_gained, falling back to generic regional boundaries. |
| Non-compliant HGVS.p & 3' Shifting | 224 | vep's 3' shifting algorithm can push in-frame deletions over the terminal stop codon, yielding stop_lost annotations and HGVS.p strings that violate strict nomenclature rules. mehari maintains the stop and produces compliant HGVS. |
| ALT == N Interpretation | 36 | mehari strictly follows the VCF specification (≥ v4.1), treating an N alternate allele as equivalent to the reference (synonymous_variant). vep treats the base as an uncharacterized alteration (protein_altering_variant). |
| Selenoprotein Support | 15 | vep lacks native selenoprotein support, resulting in inaccurate missense or stop_gained predictions. mehari natively implements the selenocysteine_gain and selenocysteine_loss SO terms. |
| Transcript Ablation Fragmentation | 7 | For deletions removing an entire transcript, mehari summarizes the event using the transcript_ablation term. vep sometimes fragments the event into constituent parts without assigning transcript_ablation. |

### HGVS Notation Concordance

`mehari` utilizes the `hgvs-rs` library, a direct Rust implementation of the established Python `biocommons-hgvs`[7] package. Because of this shared foundation, concordance between the two frameworks is structurally guaranteed to be exceptionally high. Consequently, `hgvs-rs` effectively serves as a drop-in replacement while offering a massive improvement in execution speed. In our benchmarks, `mehari` required approximately 468 CPU seconds to perform both HGVS projection and functional consequence prediction (131 µs per variant). In contrast, the Python-based `biocommons-hgvs` required roughly 104 208 CPU seconds to perform HGVS projection alone (29 169 µs per variant).

During validation on the ClinVar dataset, `hgvs-rs` reliably reproduced the expected `hgvs.c` and `hgvs.p` strings generated by its Python counterpart. Across the evaluation dataset, only 30 484 differences in `hgvs.c` or `hgvs.p` notations were observed, leading to a concordance of 99.147 %.

These remaining discrepancies are expected and stem from two systematic improvements in `mehari`’s implementation and data handling:

- Intronic Normalization: Unlike the biocommons implementation, `hgvs-rs` actively normalizes intronic variants whenever possible (but never crossing intron-exon-boundaries), leading to systematic, structurally valid divergences in notation.
- Transcript Model Recovery: While both tools utilize the same Ensembl GRCh38 cdot source data, `mehari`’s custom transcript database incorporates systematic curation. Most notably, `mehari` recovers and retains 3-prime end truncated transcripts that are otherwise dropped or handled differently.

For a full table of these discordant notations, see supplement.

### Handling Biological Edge Cases and the Sequence Ontology

Both `mehari` and `vep` classify the functional impacts of variants using the Sequence Ontology. The SO provides a standardized vocabulary graph for describing biological sequences and mutations. Crucially, both tools exclusively utilize official SO terms to represent variant consequences. This ensures strict adherence to standard nomenclature and avoids the introduction of tool-specific terms.

However, accurate variant annotation requires correctly handling complex biological edge cases that standard translation models often overlook. A primary example is the translation of selenoproteins. Under specific structural sequence contexts, the canonical `UGA` stop codon is recoded to incorporate selenocysteine (`Sec`). Legacy annotators often strictly interpret `UGA` as a stop codon, leading to erroneous `stop_gained` or missense predictions. `mehari` natively models these recoding events. It assigns accurate terms like `selenocysteine_gain` and `selenocysteine_loss` for known selenoprotein contexts rather than defaulting to generic translation tables.

In cases where the existing ontology lacked the specificity needed to describe highly specialized biological events, we actively contributed to the ontology’s expansion. For instance, we proposed adding the `selenocysteine_gain` term (SO:0002194) to accurately model mutations affecting selenoproteins^4^. Further hierarchical refinements regarding parent/child relationships for some intronic terms are currently under review^5^.

## Discussion

### The Value of “HGVS-First”

The “HGVS-first” approach enforces nomenclature normalization and hierarchical projection before evaluating functional consequences. This order of operations eliminates the ambiguity frequently seen in legacy tools. If a transcript’s coding sequence dictates a specific HGVS notation, the resulting Sequence Ontology term is deterministically bound to that logic. This tight synchronization ensures strict consistency for clinical reporting and large-scale database curation. Consequently, `mehari` serves not just as a primary annotator, but also as an efficient quality assurance tool for auditing other normalizers. To note, `vep`’s default configuration *will* lead to the HGVS descriptions being out-of-sync with the annotation, because HGVS descriptions have to be 3-prime shifted (to be valid HGVS notations), but the consequence prediction is done on the original (non-shifted) variant.

Our evaluation of edge cases between standardized terminology and biological effects allowed us to identify and improve selenoprotein annotation. This includes the integration of the `selenocysteine_gain` term (into the Sequence Ontology), which better models the special translation mechanisms of selenoproteins.

### Limitations and Future Directions

Since `mehari`’s main focus has been human clinical research, most effort has been put into building robust transcript databases for human reference assemblies (GRCh37 and GRCh38) and RefSeq and Ensembl annotations and verifying the annotations. Subsequently, the `mehari-data-tx` workflow only builds databases for the combinations of {GRCh37, GRCh38} × {Ensembl, RefSeq} plus the Ensembl and RefSeq databases merged into one. It is possible to build transcript databases by providing a standard GFF3 annotation and a FASTA file containing (n)cDNA sequences for the respective transcripts.

At the time of writing, `mehari` does not natively support deep regulatory region annotations outside the immediate transcript neighborhood. Future development will expand on these capabilities.

There is experimental support for multi-variant consequence prediction (grouping [phased] variants on the same haplotype and evaluating the compound effect on the protein sequence).

Also `mehari`’s Python bindings provide the convenient option to integrate high-speed variant effect prediction directly into larger data science ecosystems. The bindings support SPDI (`chr1:1234:A:C`), kwargs (`chromosome=“chr1”, position=1234, reference=“A”, alternative=“C”`) as well as polars dataframes with corresponding columns as input.

## Supporting information

discordant consequence cases (vep)

discordant hgvs cases (biocommons-hgvs)

## Availability

`mehari` is open-source software distributed under the MIT License. We strive to provide best-effort support via GitHub issues.

The source code and pre-built binaries are available across several channels:

- Source code: https://github.com/varfish-org/mehari
- Binary: Available via bioconda (`conda install bioconda::mehari`)
- Python bindings: Available via PyPI (`pip install mehari`) and bioconda (`conda install bioconda::mehari-python`)
- snakemake-wrappers
- Database workflow and pre-built databases: https://github.com/varfish-org/mehari-data-tx
- Website: https://varfish-org.github.io/mehari

## Footnotes

1 This comparison is a case of comparing apples and oranges: while the 24GB vep cache does provide more information like regulatory motifs and pre-calculated SIFT/PolyPhen matrices, its footprint is inflated by the way the information is serialized, whereas mehari’s modular architecture prioritizes clinical relevance and data density through optimized binary protobuf and RocksDB formats.

2 This mode enables heuristics designed to mimic vep’s output, such as making certain SO terms less specific (e.g., using `inframe_deletion` instead of `disruptive_inframe_deletion`) and explicitly suppressing certain overlapping terms.

3 The exact output of vep depends heavily on the combination of CLI options used. While explicitly specifying both `--shift_hgvs 1` and `--shift_3prime 1` results in an error, specifying only `--shift_3prime 1` seems to incorporate the effects of `--shift_hgvs 1` implicitly. We utilized this configuration because it produced the closest match for both correct HGVS notations and sequence consequences.

4 https://github.com/The-Sequence-Ontology/SO-Ontologies/issues/653

5 https://github.com/The-Sequence-Ontology/SO-Ontologies/issues/654

## Bibliography

[1] F. Mölder et al., “Sustainable data analysis with Snakemake,” F1000Research, vol. 10, p. 33, 2021.

[2] W. McLaren et al., “The ensembl variant effect predictor,” Genome biology, vol. 17, pp. 1–14, 2016.

[3] K. Wang, M. Li, and H. Hakonarson, “ANNOVAR: functional annotation of genetic variants from high-throughput sequencing data,” Nucleic acids research, vol. 38, no. 16, p. e164–e164, 2010.

[4] M. Stromberg, R. Roy, J. Lajugie, Y. Jiang, H. Li, and E. Margulies, “Nirvana: clinical grade variant annotator,” in Proceedings of the 8th ACM International Conference on Bioinformatics, Computational Biology, and Health Informatics, 2017, p. 596.

[5] P. Cingolani et al., “A program for annotating and predicting the effects of single nucleotide polymorphisms, SnpEff: SNPs in the genome of Drosophila melanogaster strain w1118; iso-2; iso-3,” fly, vol. 6, no. 2, pp. 80–92, 2012.

[6] T. Hartmann, C. Schröder, E. Kuthe, D. Lähnemann, and J. Köster, “Insane in the vembrane: filtering and transforming VCF/BCF files,” Bioinformatics, vol. 39, no. 1, p. btac810, 2023.

[7] M. Wang et al., “hgvs: A Python package for manipulating sequence variants using HGVS nomenclature: 2018 Update,” Human mutation, vol. 39, no. 12, pp. 1803–1813, 2018.

